# Robust and scalable manifold learning via landmark diffusion for long-term medical signal processing

**DOI:** 10.1101/2020.05.31.126649

**Authors:** Chao Shen, Yu-Ting Lin, Hau-Tieng Wu

## Abstract

Motivated by analyzing long-term physiological time series, we design a robust and scalable spectral embedding algorithm, coined the algorithm RObust and Scalable Embedding via LANdmark Diffusion (ROSE-LAND). The key is designing a diffusion process on the dataset, where the diffusion is forced to interchange on a small subset called the *landmark set*. In addition to demonstrating its application to spectral clustering and image segmentation, the algorithm is applied to study the long-term arterial blood pressure waveform dynamics during a liver transplant operation lasting for 12 hours long.

## 1. Introduction

Learning from data has been an intriguing topic in many scientific fields, particularly the biomedical field. A learning procedure is taking existing data as the “knowledge” or “experience” to interpret incoming data and help field experts make decision. In the clinical medicine, it has been widely debated that eventually a well trained artificial intelligence (AI) by a proper learning procedure could replace physicians. While it is not impossible, we believe that there is still a long way to go. No matter how, a general consensus is that such a system would augment physicians, and release physicians’ precious time by automatizing some routine and time-consuming workflow so that physicians.

Motivated by its importance, various datasets of different data types from clinics have been extensively explored from this learning perspective, like electronic health record [30], medical imaging [17], genomic data [38], biomedical waveforms (or time series, signals) [15], etc. To our knowledge, however, how to extract intrinsic dynamics underlying biomedical waveforms for clinical usage is relatively less discussed, and most of existing literature focus on simplifying the waveform information into few scalars [7, 34, 2, 45]. While this has been successfully applied to clinical medicine, we may loss information encoded in the original waveform. One solution to depict intrinsic dynamics directly from the original waveform is obtaining as many features as possible, and selecting suitable parameters for the learning purpose [19]. Another solution is applying manifold learning algorithms to the *original* physiological waveforms [28, 48]. The basic idea in [28, 48] is truncating the physiological waveform into pieces according to some rules, and then apply the spectral embedding algorithm, like the diffusion maps (DM) [9], to embed those pieces into a finite dimensional Euclidean space, which represents the intrinsic dynamics. If the physiological waveform is embedded into the three dimensional Euclidean space, the physiological waveform is converted into a three dimensional image so that users can visualize the waveform from a different perspective.

It is commonly believed that more data lead to more comprehensive “knowledge” in the learning procedure, and hence more informative features and better visualization of the intrinsic dynamics. We thus hypothesize that if the technique shown in [28, 48] could be applied to analyze long-term physiological waveforms of length on the order of days or weeks, it will be beneficial to supporting the short-term memory of the human brain when handling long-term physiological waveforms. Specifically, due to the short-term memory limitation, it is easy to overlook information hidden in the long-term waveform; for example, what is the relationship between the waveform in the first hour of the operator and the 7th hour? Will the surgeon remember the useful information in the first hour after 5 hours? While this idea is natural, unfortunately, it is prohibited by the computational complexity inherited in most spectral decomposition based machine learning algorithm, including the DM applied in [48].

Recall that the DM is based on the eigendecomposition of the graph Laplacian (GL) matrix. The algorithm has been shown to perform well when the database is “tiny”, like in the order of 10^3^ ∼ 10^4^. However, when the database gets larger, like in the order of 10^6^ or above, the algorithm is challenged by the scalability issue. Take electrocardiogram (ECG) into account. There are 10^3^ ∼ 10^4^ cycles in 1 hour long ECG, and roughly 10^6^ cycles in 14 days. In practice, it is natural to consider subsampling the dataset, however, we may loss information. As a result, although the DM works well and provides alternative clinical information [48], it is limited to datasets of length about one hour.

There have been several solutions toward this scalability challenge. One usual technique is the k-nearest neighbor (kNN) scheme. However, it is not robust to noise. Specifically, when the dataset is noisy and the neighboring information is *not* provided, obtaining a reliable kNN information is challenging. A randomized kNN approach is recently considered in [29]. Another practical solution is directly subsampling the dataset, and then recovering the information of interest by the *Nyström extension* [10]. This approach is also called the *Nyström low-rank approximation* [6], the *kernel extension method* [14], or in general the *interpolative decomposition* [32]. This approach has several theoretical backups, for example [6], and has been widely applied. While it works well for some missions, this approach is limited by the information loss during the subsampling process. Yet another approach is speeding up the matrix decomposition by randomization. For example, we can speed up the algorithm by constructing a thin matrix by taking a random subset of columns of the GL matrix and evaluating the singular value decomposition (SVD) [32]. While this approach has been widely applied, to the best of our knowledge, we have limited knowledge about how it helps the spectral embedding algorithms, and how robust it is to the inevitable noise.

Unlike the above, in this paper we propose a novel algorithm that resolves two common challenges when we apply spectral embedding algorithms—robustness and scalability. The algorithm is intuitive and can be summarized in three steps. First, we find a “small” subset of points from the whole dataset, either randomly or by design, or collect a separate clean point cloud of small size, which we call a *landmark set*. Second, we construct an affinity matrix recording the affinities between points in the whole dataset and the landmark set, and normalize it properly. This normalized affinity matrix is thin; that is, there are fewer columns than rows. Third, evaluate the singular vectors and singular values of the normalized affinity matrix, and embed the dataset using the singular vectors and singular values. As we will make clear soon, this algorithm is directly related to the diffusion process. We coin the proposed algorithm the *RObust and Scalable Embedding via LANdmark Diffusion* (Roseland). The solution is generic and not limited to analyze the motivating physiological waveform problem.

In addition to demonstrating its application to spectral clustering and image segmentation, the Roseland is applied to study the long-term arterial blood pressure waveform dynamics during a liver transplant operation lasting for 12 hours long. For the sake of self-containedness, we also summarize and explain the theoretical results shown in [40], particularly the spectral convergence and robustness of Roseland under the manifold setup.

## 2. The Proposed Roseland Algorithm

We assume that we have a data set 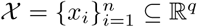. Take a set 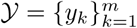, which might or might not be a subset of 𝒳. We call 𝒴 the *landmark set*. Fix a non-negative kernel function *K* : ℝ_≥0_ → ℝ_+_ with proper decay and regularity; for example, a Gaussian function.

First we construct a *landmark-set affinity matrix W* ^(r)^ ∈ ℝ^*n*×*m*^, which is defined as

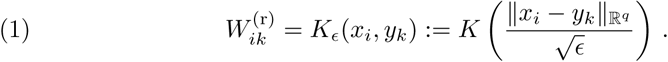

That is, the (*i, k*)-th entry of *W*^(r)^ is the similarity between the *i*-th data point and *j*-th landmark, and clearly the larger the distance between two points (or the two points are less similar), the smaller the similarity. Next compute a diagonal matrix *D*^(R)^ as

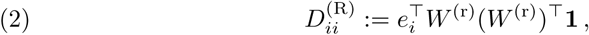

where **1** is a *n*×1 vector with all entries 1, and *e*_*i*_ is the unit vector with 1 in the *i*-th entry. 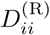 is called the degree of the *i*-th data point *x*_*i*_. Intuitively, it represents how strong *x*_*i*_ is attached to the data set. With *W* ^(r)^ and *D*^(R)^, we evaluate the SVD of (*D*^(R)^)^−1*/*2^*W*^(r)^:

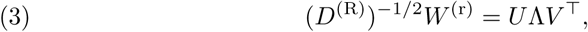

where the singular values *σ*_1_ ≥ *σ*_2_ ≥ … ≥ *σ*_*m*_ ≥ 0 are on the diagonal of the diagonal matrix Λ. Set *Ū* := (*D*^(R)^)^−1*/*2^*U*. Take *q*′ ∈ ℕ so that *q*′ ≤ *m*. Let *Ū*_*q*′_∈ ℝ^*n*×*q*′^ to be a matrix consisting of the second to the (*q*′ + 1)-th columns of *Ū* and 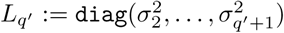. Finally we define the Roseland embedding as

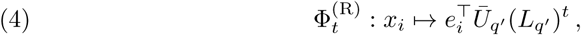

where *t* > 0 is the chosen diffusion time, in other words, the *i*-th data point *x*_*i*_ is embedded using the *i*-th row of *Ū*_*q*′_ entry-wisly rescaled by 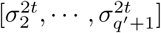. See Algorithm 1 for a summarization of the Roseland algorithm. We thus define the associated *Roseland diffusion distance* (RDD) by

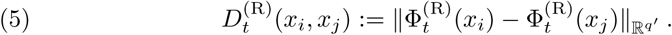

### Algorithm 1 The pseudo-code of Roseland.

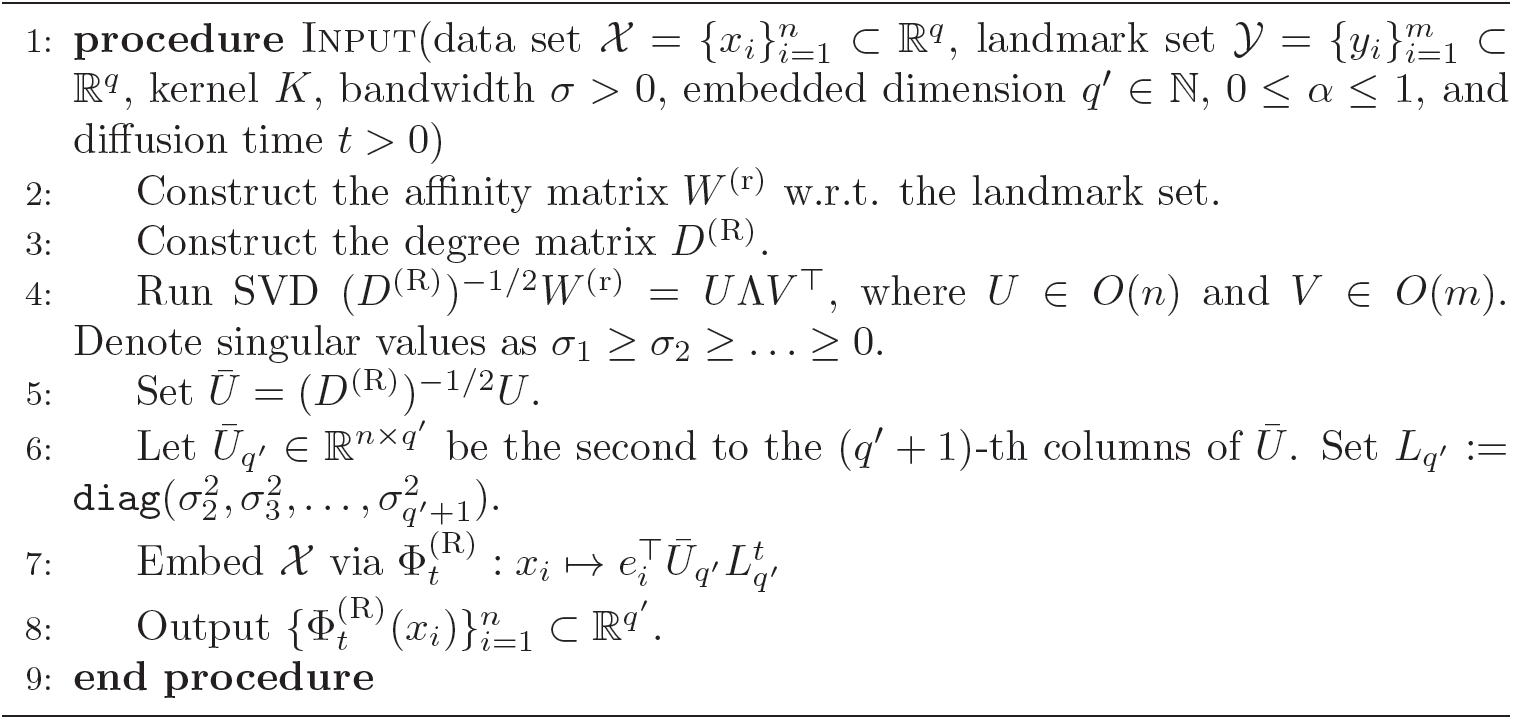

Note that the Roseland induces a new affinity matrix on the data set 𝒳 via

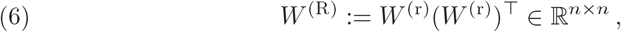

where *W* ^(r)^ is the landmark-set affinity matrix (1). We call *W* ^(R)^ the *landmark-affinity matrix*, which is positive and positive-definite. We remark that traditional affinity matrices between data points are often constructed from *one* global pre-fixed kernel *K*, while in Roseland we cannot find a global fixed kernel 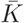 and a bandwidth 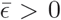 so that 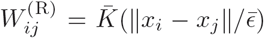 for all *i, j* in general. Hence Roseland can provide us with more dynamic similarity measurements between data points based on the landmarks. Also note that *A*^(R)^ := (*D*^(R)^)^−1^*W* ^(R)^ is a transition matrix on 𝒳, which means each of the row in *A*^(R)^ sums to 1. This is because *D*^(R)^ is the degree matrix associated with the landmark-affinity matrix *W* ^(R)^ by construction. Hence, we can view the similarity measure between data points *x*_*i*_ and *x*_*j*_ as a Markov or diffusion process through the landmarks.

### 2.1. Reference set as subset of the data

In cases that we may not be able to acquire additional data points as landmark set but have to select the landmark set from the available dataset, we propose to first sample *m* landmarks, denoted by 𝒴, from 𝒳 so that 𝒴 is independent of 𝒳 *\* 𝒴. Then, we apply the Roseland on 𝒳 *\* 𝒴 using 𝒴 as landmarks, and extend the embedding to 𝒴 by the Nyström extension. When |𝒴| ≪ |𝒳 |, the discrepancy of this approach and the independence setup with original Roseland is negligible, and will asymptotically vanish.

### 2.2. Related methods

#### 2.2.1. Graph Laplacian and Diffusion Maps

We now review the GL and compare Roseland with the well-known algorithm diffusion map (DM) [9]. First, pre-fix a kernel function *K* and a bandwidth parameter *ϵ* > 0. Then, compute the affinity matrix *W* ∈ ℝ^*n*×*n*^ by

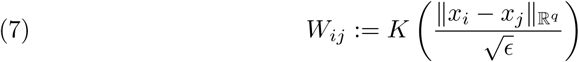

and the corresponding *degree matrix D* ∈ ℝ^*n*×*n*^, which is a diagonal matrix defined as 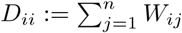. For a fixed *α* ∈ [0, 1], the *α-normalized* affinity matrix *W* ^(*α*)^ ∈ ℝ^*n*×*n*^ [9] is defined as 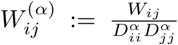, where 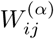 is called the *α-normalized affinity* between *x*_*i*_ and *x*_*j*_. Note that *W* ^(0)^ = *W* defined in (7). In some applications when we want to remove the density effect caused by data sampling, we set *α* = 1. With the *α*-normalized affinity matrix *W* ^(*α*)^, one can analogously define the associated degree matrix *D*^(*α*)^ ∈ ℝ^*n*×*n*^ by 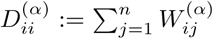. The GL is defined as *L*^(*α*)^ := *I* − *A*^(*α*)^, where

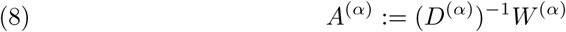

is the associated transition matrix. Clearly, *A*^(*α*)^ is row stochastic, and it defines a random walk on the dataset 𝒳. We mention that the transition matrix (*D*^(R)^)^−1^*W* ^(R)^ in the Roseland algorithm can be viewed as an alternative way of constructing a Markov process on the dataset 𝒳. Since *A*^(*α*)^ is similar to (*D*^(*α*)^)^−1*/*2^*W* ^(*α*)^(*D*^(*α*)^)^−1*/*2^ we can find its eigendecomposition with eigenvalues 1 = λ_1_ > λ_2_ ≥ … ≥ λ_*n*_ and the associated eigenvectors *ϕ*_1_, …, *ϕ*_*n*_. Denote *ϕ*_*i*_ the *i*-th right eigenvector of *A*^(*α*)^. Recall in Roseland, we perform the SVD decomposition in (3), which is a parallel step of the eigen-decomposition of *Ā*^(0)^.

Among various algorithms, we focus on the well-known DM algorithm that we shall see to be closely related to Roseland. With the spectral decomposition of the GL, the chosen normalization *α*, embedding dimension *q*′ and *diffusion time t*, the DM embeds 𝒳 via the map

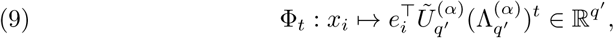

where 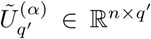 to be a matrix consisting of the second to the (*q*′ + 1)-th columns of *Ũ*^(*α*)^ and 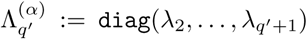. The diffusion distance (DD) with the diffusion time *t* > 0 is defined as

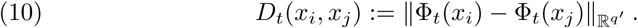

Recall the Roseland embedding (4) and the RDD (5) to notice the close relation between the DM embedding and the DD.

As it is expensive to perform eigen-decomposition of dense matrices, one common practice of DM or general spectral embedding methods is to use the kNN scheme to construct a rather sparse affinity matrix; that is, set *W*_*ij*_ = 0 when *x*_*j*_ is not within the first *k* nearest neighbors of *x*_*i*_, where *k* is chosen by the user. Another way is to use a compactly supported kernel *K*. For example, *K*(*t*) is 1 when *t* ∈ [0, 1] and 0 when *t* > 1.

#### 2.2.2. Nyström Extension

Another directly related algorithm is the common Nyström extension [3, 14, 50]. The basic idea is running the eigen-decompostion on a small subset of the whole database, and then extending the eigenvectors to the whole dataset. In this work, we apply the modified Nyström extension [22, 43] that is suitable for the diffusion process. See [40] for a summary of this algorithm.

## 3. Theoretical Results

In this section, we summarize two theoretical results shown in the accompanying paper [40], one is the asymptotic spectral convergence and one is the robustness of Roseland under the manifold setup.

### 3.1. Manifold model

Denote our observed dataset by 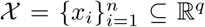. Assume is independently and identically (i.i.d.) sampled from a random vector *X* : (Ω, ℱ, 𝕡) → ℝ^*q*^, where the range of *X* is assumed to be supported on a *d*-dimensional compact smooth Riemannian manifold (*M*^*d*^, *g*) without boundary isometrically embedded in ℝ^*q*^ via *ι* : *M*^*d*^ ↪ ℝ^*q*^. Suppose the density function of *X* on *M* is denoted as *p*_*X*_. Similarly, denote the landmark set by 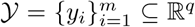. Assume 𝒴 is i.i.d. sampled from a random vector *Y* : (Ω, ℱ, 𝕡) → ℝ^*q*^, where the range of *Y* is assumed to be (*M*^*d*^, *g*) as well. Suppose the density function of *Y* on *M* is denoted as *p*_*Y*_. Furthermore, assume that *X* and *Y* are independent.

### 3.2. Spectral convergence

Let {*v*_*n*_}_*n*∈ℕ_ be the set of eigenvectors of the matrix (*D*^(R)^)^−1^*W* ^(R)^ evaluated from the dataset 𝒳 and the landmark 𝒴. It is shown in [40] that the eigenvectors {*v*_*n*_}_*n*∈ℕ_ asymptotically converge to the eigenfunctions of the Laplace-Beltrami operator of (*M, g*) as *n* → ∞. Hence, by the well established spectral geometry theory [4, 36], the Roseland embedding recover the underlying manifold.

First, note that the eigenvectors *v*_*n*_ are in difference Euclidean spaces for different *n*, hence they cannot be compared directly with the eigenfunctions of the Laplace-Beltrami operator, which are smooth functions on *M*. To make sense of the comparison, we find a sequence of functions *f*_*n*_ ∈ *C*(*M*), such that the restriction of *f*_*n*_ on the data 𝒳 is *v*_*n*_; that is, *f*_*n*_(*x*_*i*_) = *v*_*n*_(*i*), for *i* = 1, …, *n*. After that, we study the convergence of {*f*_*n*_} as *n* → ∞. We start with some definitions.

#### Definition 1.

*Take* 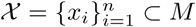. *Define the following functions*

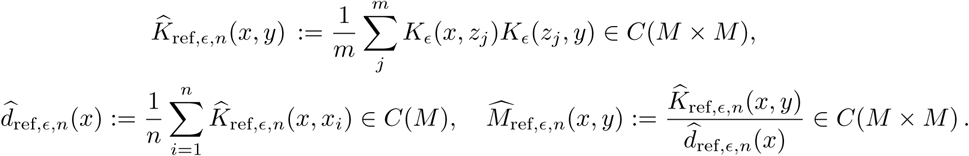

*Let*

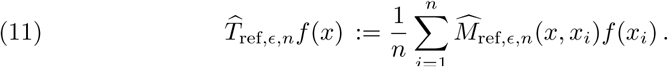

*Moreover, define the restriction operator ρ*_*n*_ : *C*(*M*) → ℝ^*n*^ *by ρ*_*n*_ : *f* ↦ [*f* (*x*_1_), *f* (*x*_2_), …, *f* (*x*_*n*_)]^T^.

The following Lemma essentially says that if we restrict the eigenfunctions of 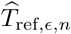 on the data, we obtain the eigenvectors of (*D*^(R)^)^−1^*W* ^(R)^.

#### Proposition 1.

[40] *Let U*_*n*_ := (*D*^(R)^)^−1^*W* ^(R)^, *then* 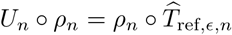. *More-over, we have the following one to one correspondence.*

1. *If f* ∈ *C*(*M*) *is an eigenfunction of* 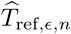 *with the eigenvalue* λ, *then the vector v* := *ρ*_*n*_*f is an eigenvector of U*_*n*_ *with the eigenvalue* λ. *Moreover, suppose* λ ≠ 0 *is an eigenvalue of* 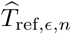 *with the eigenfunction f. If we let v* := *ρ*_*n*_*f, then f satisfies*

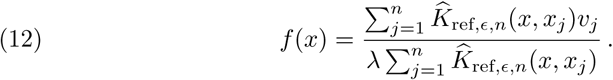
2. *If v is an eigenvector of U*_*n*_ *with the eigenvalue* λ ≠ 0, *then f defined in* (12) *is an eigenfunction of* 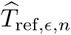 *with the eigenvalue* λ.

Denoted by Δ the Laplace-Beltrami operator on (*M, g*). Let (λ_*i*_, *u*_*i*_) be the *i*-th eigenpair of −Δ, where the eigenvalues λ_*i*_ are in the ascending order. Under our manifold setup and by the well known elliptic theory, the spectrum of −Δ is discrete ∞with as the only accumulation point, and each eigenspace is of finite dimension. Correspondingly, let (λ_*ϵ,n,i*_, *u*_*ϵ,n,i*_) be the *i*-th eigenpair of 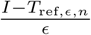, where the eigenvalues λ_*ϵ,n,i*_ are in the ascending order as well. We assume that ‖*u*_*i*_‖_2_ = ‖*u*_*ϵ,n,i*_‖_2_ = 1. With the above preparation, we have the following theorem.

#### Theorem 1

(Spectral convergence). [40] *Fix K* ∈ ℕ. *Suppose the kernel is Gaussian; that is*, 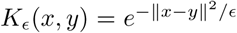. *Suppose* λ_*i*_ *is simple. Suppose m* = *n*^*β*^, *where β* ∈ (0, 1), *ϵ* = *ϵ*(*n*) *so that ϵ* → 0 *and* 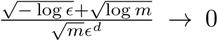, *as n* → ∞, *and* 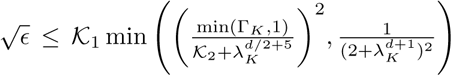, *where* Γ_*K*_, *𝒦*_1_ *and 𝒦*_2_ > 1 *are introduced in* [40]. *Then, when p*_*Y*_ *is properly chosen so that* 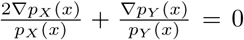, *there exists a sequence of signs a*_*n*_ *such that with probability* 1 − 𝒪 (*m*^−2^), *for all i* < *K, we have*

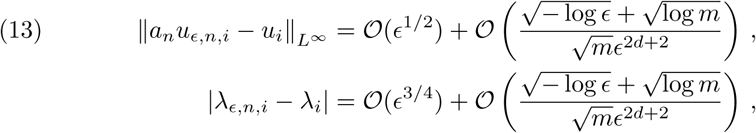

*where the implied constants depend on the kernel, the curvature of M, p*_*X*_ *and p*_*Y*_

We see that the error term converges to zero if we further assume that 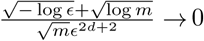. We mention that when λ_*i*_ is not simple, the theorem still holds, but we need to use the eigenprojection to describe the eigenvector convergence statement.

### 3.3. Robustness

In real applications, the dataset is often corrupted by noise. In this case, spectral embedding algorithms might lead us to a low quality, or even misleading result. This robustness issue for DM was studied in [11, 12] and some solutions are proposed. It is suggested in [12] to work with the complete graph or a graph with a large number of kNN, and force the random walk non-lazy. While this solution works, however, it is not scalable. Roseland, on the other hand, automatically enjoys the desired robustness property. Note that we measure similarities between data points by diffusing through *all* landmarks in Roseland. This seemingly simple step has a significant consequence. It can be viewed as a surrogate of knowing *true neighbors* in the kNN scheme, and it explains the robustness of Roseland. To appreciate this significance, recall that when the neighboring information is not provided and if the kNN approach is considered, we need to estimate the neighbors. However, estimating neighbors from noisy data is error-prone and wrong neighboring information corrupts the DM.

To show the robustness of Roseland, on top of the manifold model in Section 3.1, we assume that the data set and the landmark set are corrupted by additive ambient space noise. That is, we observe 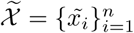 and 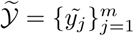:

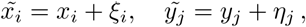

where *ξ*_*i*_ and *η*_*j*_ are noise. We assume that *x*_*i*_, *y*_*j*_, *ξ*_*i*_ and *η*_*j*_ are independent. For more practically challenging purpose, we consider large *q* large *n* setup, that is, the ambient space dimension grows asymptotically as the dataset size *n*. To simplify the discussion, we focus on Gaussian noise, and refer readers with interest to [40] for more complicated non-Gaussian noise model and results.

#### Assumption 3.1

(Gaussian noise model). *Suppose the noise contaminating the dataset is ξ*_*i*_ ∼ 𝒩 (0, Σ) *with mean* 0 *and covariance* Σ *and the noise contaminating the landmark set is* 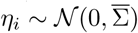 *with mean* 0 *and covariance* 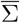. *Assume ξ*_*i*_ *and η*_*j*_ *are independent. Suppose* 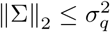 *and* 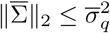, *where σ*_*q*_ ≥ 0 *and* 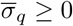.

Recall the Roseland algorithm in Section 2. Denote the Roseland embedding with diffusion time *t* from clean data and noisy data by Φ*L*^*t*^ ⊂ ℝ^*n*×*q*′^ and 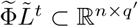 respectively. We analyze the discrepancy between these two embeddings. Since the embeddings are free up to rotations and reflections, we quantify 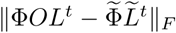, where *O* ℝ^*q*′ ×*q*′^ is some orthogonal matrix that aligns the embeddings, and ‖·‖_*F*_ is the Frobenius norm. We have the following theorem.

#### Theorem 2

(Robustness of Roseland). [40] *Assume the point clouds* 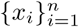 *and* 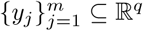 *are i.i.d. sampled from the underlying manifold. Let* 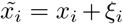 *and* 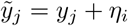, *where the noises ξ*_*i*_ *and y*_*j*_ *satisfy Assumption 3.1 are independent of x*_*i*_ *and y*_*j*_. *Also, assume*

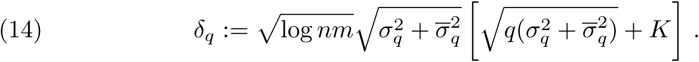

*Fix q*′ ∈ ℕ. *According to Theorem 1, pick a sufficiently small ϵ* = *ϵ*(*q*′) > 0 *so that the first q*′ *non-trivial singular values are sufficiently away from zero when n is sufficiently large. Denote W* ^(r)^ *and* 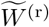 *to be the landmark-set affinity matrices from clean and noisy datasets respectively. Denote* Φ*L*^*t*^ ∈ ℝ^*n*×*q*′^ *and* 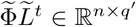 *to be Roseland embeddings from W* ^(r)^ *and* 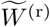 *respectively. Then, for fixed t* ≥ 0 *and q*′ ∈ ℕ, *we have*

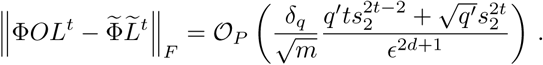

*where O* ℝ^*q*′×*q*′^ *is an orthogonal matrix, and s*_2_ *are the largest non-trivial singular value of Roseland from the clean.*

The Theorem says that the bandwidth *ϵ* should chose “large” enough so that the noise impact on the embedding is alleviated. This fulfills the intuition that we can tame the noise by aggregating more independent noise, since a larger bandwidth indicates including more noisy data locally. Note that while *δ*_*q*_ → 0 when *n* → ∞, it does not mean that the noise impact is small when *n* → ∞. We shall quantify the noise impact by 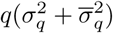 in *δ*_*q*_, which can be viewed as the *total energy of noise* in the data. From this perspective, we mention that Roseland can tolerate large noise, with the noise level up to *σ*_*q*_ = *q*^−(1*/*4+*a*)^ for arbitrary constant *a* > 0. In this setup, the total energy in noise blows up since 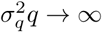. For more discussion, see [40].

Moreover, when the landmark set is noise free, we could achieve a better convergence since 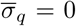. This condition is related to the situation that we are able to collect a small clean dataset as the landmark set in addition to the large but noisy dataset. This situation is commonly encountered in real life. For example, in the medical field, collecting a clean dataset of high quality is usually labor-intensive and expensive. However, it is relatively easy to collect a large dataset from a rather cheap equipment, in exchange of the data quality.

## 4. Numerical Results

In this section, we show how to apply Roseland to speed up spectral clustering and image segmentation when the dataset is big.

### 4.1. Spectral clustering

MNIST is a dataset consisting of labelled handwritten digits from 0 to 9, in the form of 28 × 28 gray-scale images [23]. There are 60, 000 training images and 10, 000 testing images. For the spectral clustering purpose, we include all 70, 000 data points. For a fair comparison between Nyström extension and Roseland, we fix the landmark set of size 250; that is *β* ≈ 0.5.

A spectral clustering algorithm consists of two steps: the first step is performing a chosen spectral embedding, and the second step is applying the *K*-mean clustering. In our experiment, we set *K* = 10. Let 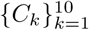 be the clusters obtained by the spectral clustering. To evaluate the performance of the spectral clustering based on different spectral embedding algorithms, including the diffusion map, Roseland and Nyström extension, we consider the following procedure. Denote the data set by 𝒳, so 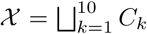. Let *f* : 𝒳 → *y* be the true predictor function and 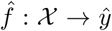 be the estimated predictor function that is constant on *C*_*k*_ for ∀*k*. Here, 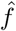 is defined in the following way:

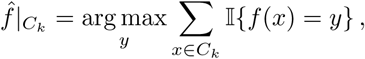

for *k* = 1, …, 10, where 𝕀 is the indicator function. In other words, for each cluster, we take a vote to decide the label it is associated with. As MNIST is almost balanced, we compute the accuracy as follows:

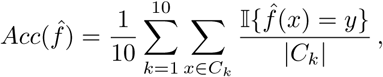

where |·| is the size of the cluster. In addition to running the above on the clean MNIST, we also run the algorithm on the MNIST with additive Gaussian noise. Specifically, we first normalized all pixel values, then entry-wisely add Gaussian noise with mean 0 and standard deviation 0.2.

We plot 3 dimensional spectral embeddings for a visual comparison. To enhance the color contrast for the visualization purpose, we only show the first 5 digits. See Figure 2 for the embedding result. In our simulations, the DM (with the sparse kernel matrix) of the whole MNIST dataset took approximately 1,500 seconds, the Nyström extension took about 5 seconds, and the Roseland took about 15 seconds. The accuracies are reported in Figure 3. All of the simulations were done on a Linux machine with 4-core 3.5Ghz i5 CPUs and 16GB memory.

**Figure 1.**
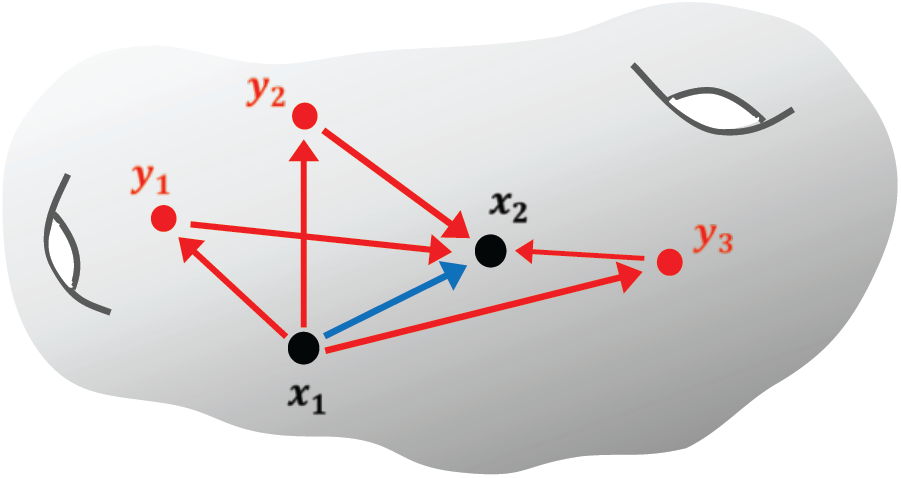
Main idea of Roseland: to measure the similarity between *x*_1_ to *x*_2_, instead of diffuse from *x*_1_ to *x*_2_ directly, we take a detour and first diffuse *x*_1_ to the landmarks *y*_1_, *y*_2_, *y*_3_, and then diffuse from the landmarks to *x*_2_.

**Figure 2.**
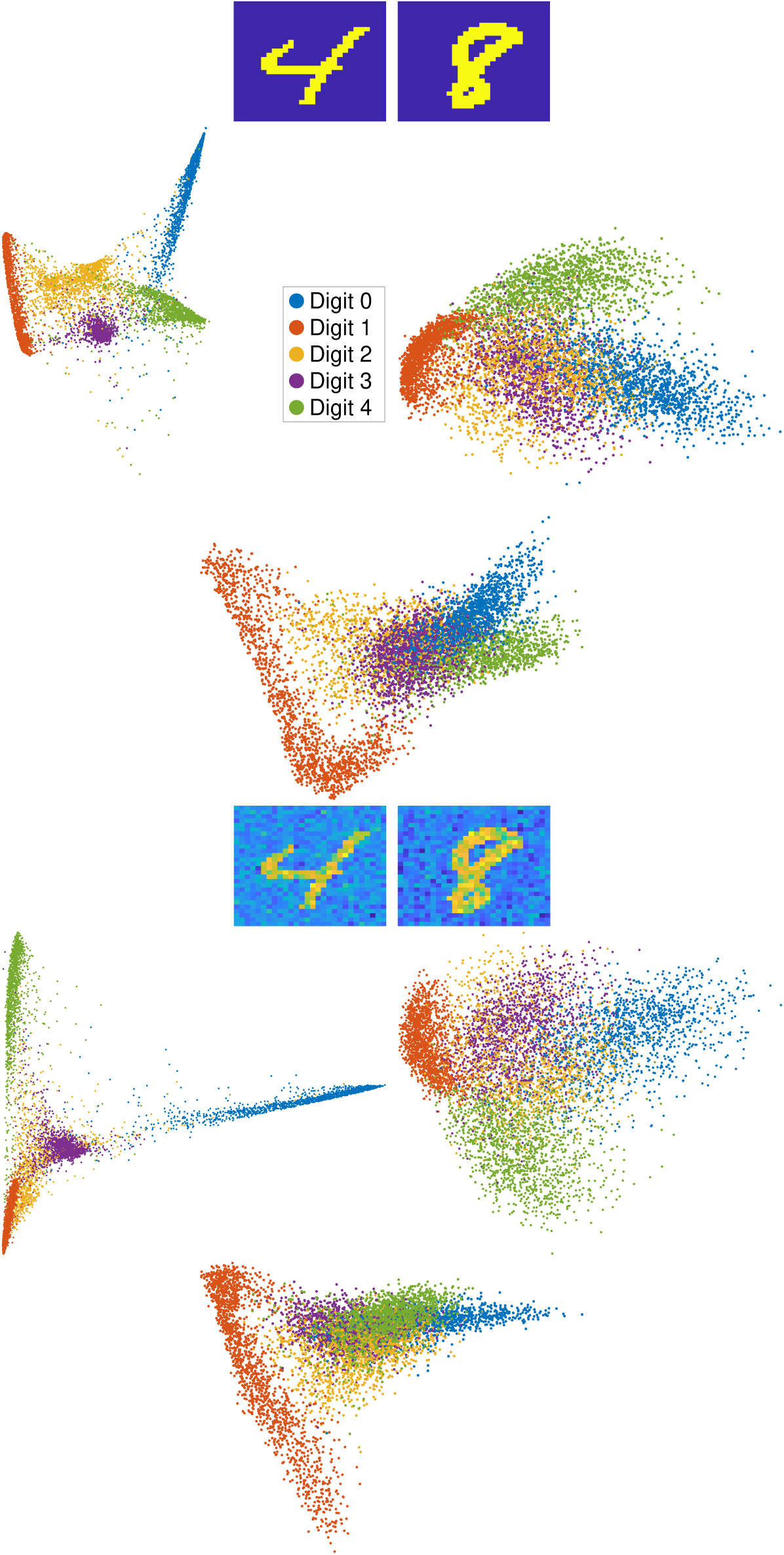
First row: Examples of clean digits. Second row: embeddings of digits 0 to 4 using clean MNIST, from left to right: the DM embedding, the Nyström extension embedding, and the Roseland embedding. Third row: Examples of noisy digits. Fourth row: embeddings of digits 0 to 4 using noisy MNIST, from left to right: the DM embedding, the Nyström extension embedding use clean landmars, and the Roseland embedding use clean landmarks.

**Figure 3.**
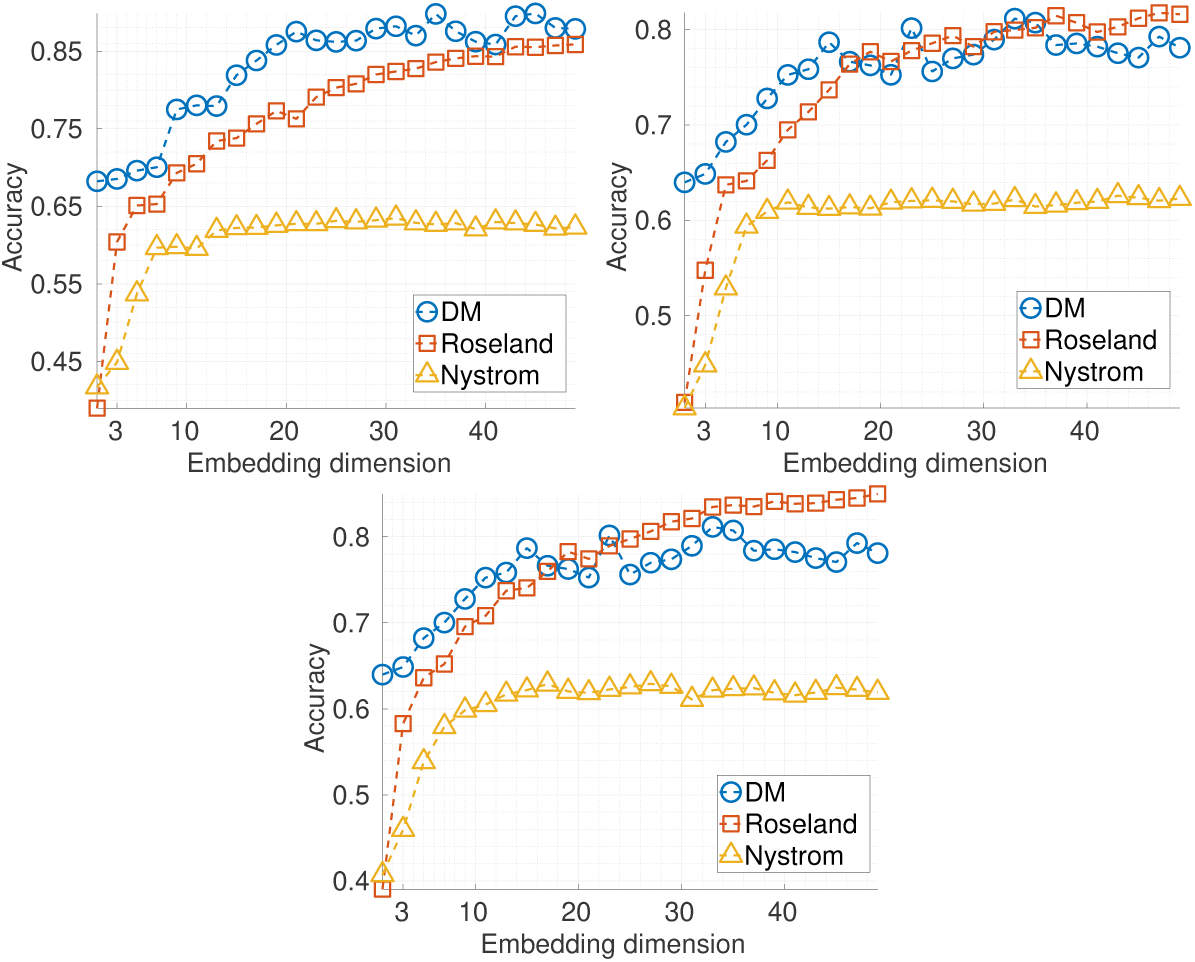
Accuracies of three spectral clustering. From left to right: clean MNIST, noisy MNIST with noisy landmarks, noisy MNIST with clean landmarks.

We mention that the two largest experiments we are aware of with the MNIST dataset are [16] and [26]. In [16], the authors improved the convergence of the kernel Hebbian algorithm to accelerate the kernel Principle Component Analysis algorithm, and the total run time is more than 50 hours on 60, 000 images. No classification accuracy was reported. The experiments were performed on an AMD Athlon 2.4 GHz CPU with 2 GB main memory and 512 kB cache. In [26], the authors applied a similar idea to HKC, but with a different normalization. Similar to [51], the algorithm output clusters rather than embeddings. The algorithm took about 100 seconds on 70, 000 MNIST images, and the overall accuracy achieved 70%. In [35], the authors adapted the Nyström idea and proposed a greedy algorithm to build the subset via projection. In their experiments, they restricted the run time to be less than 14 seconds, and computed the embeddings of 1,300 MNIST images consisting of only 0’s and 1’s with image sizes scaled down from 28 × 28 to 14 × 14.

### 4.2. Image segmentation

Next, we apply the spectral clustering algorithm to the image segmentation problem [41]. The first two images shown in Figure 4 are of size 481 × 321 from the Berkeley Segmentation Dataset and Benchmark [31] and the third image is of size 582 × 640 from the COCO data set [27]. To make the task more challenging, we corrupted the images with additive noise. Specifically, we first normalized all pixel values, then entry-wisely add Gaussian noise with mean 0 and standard deviation 0.2, see Figure 5.

**Figure 4.**
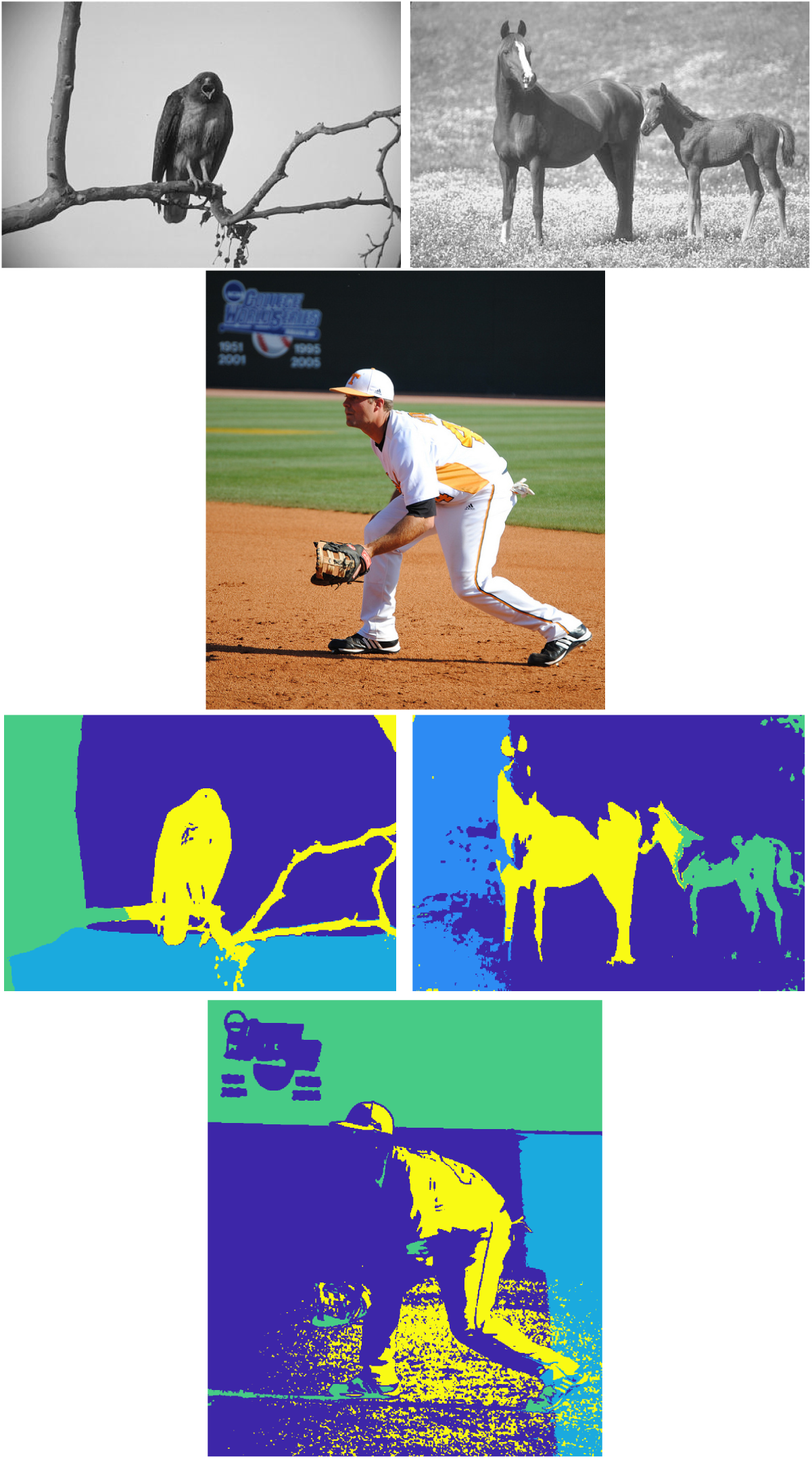
Top row: first and second images are from the Berkeley Segmentation Dataset and Benchmark, of size 481 × 321, the third one is from COCO, of size 582 × 640. Bottom row: Image segmentations by spectral clustering based on Roseland using three eigenvectors.

**Figure 5.**
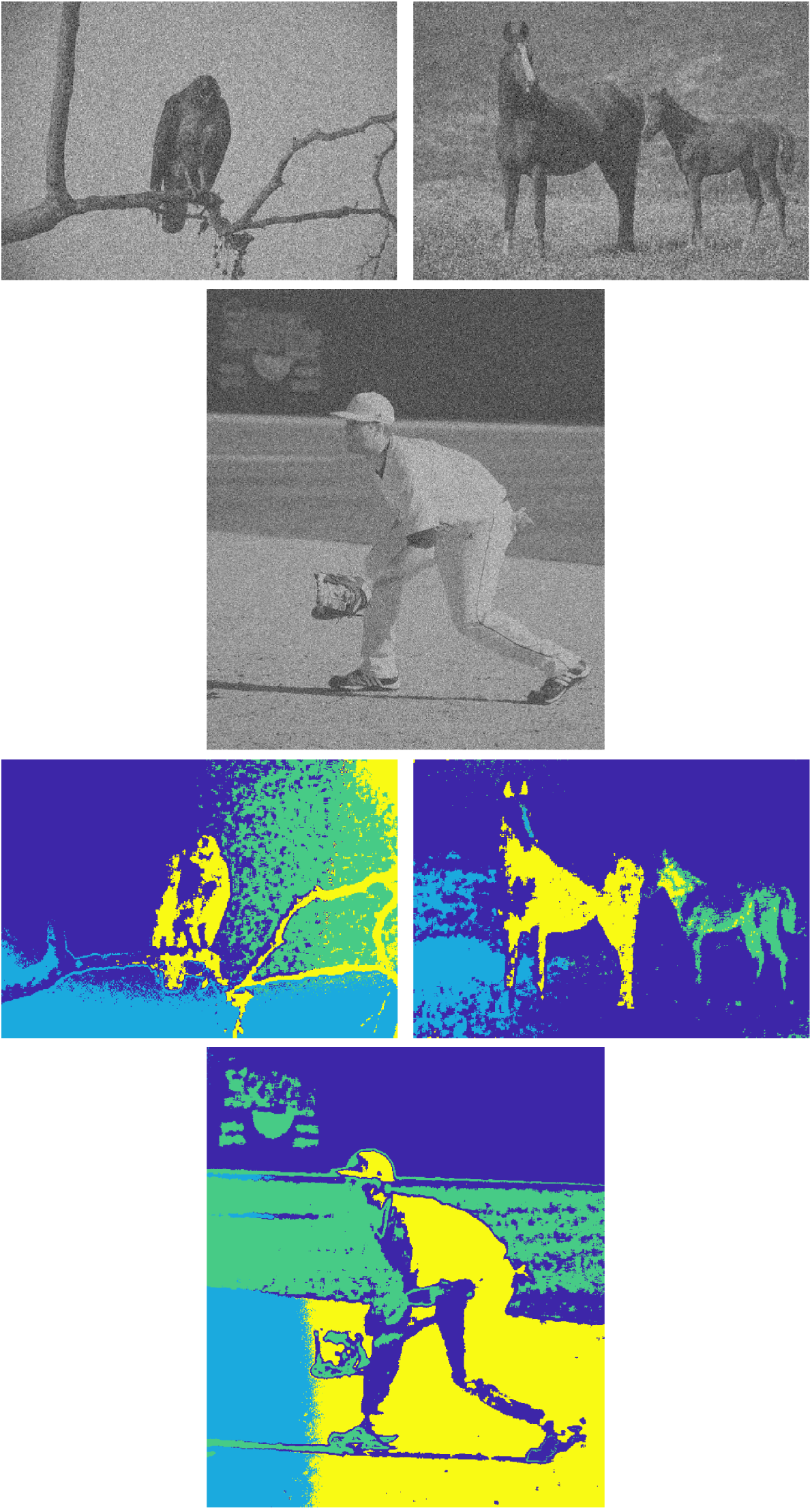
Top row: first and second images are from the Berkeley Segmentation Dataset and Benchmark, of size 481 × 321, the third one is from COCO, of size 582 × 640 corrupted by Gaussian noise. Bottom row: Image segmentations by spectral clustering based on Roseland using three eigenvectors.

Due to the data size, we only applied the Roseland to perform image segmentation. We slightly modified the algorithm proposed in [41]. Specifically, for each pixel, we attach a 3 × 3 patch surrounding the pixel. For all images, we fix the landmark set size to be 150. For the two images of size 481 × 321, the Roseland based image segmentation took about 5 seconds, and for the image of size 582 × 640, it took about 12 seconds. The results are shown in Figures 4 and 5.

## 5. Liver Transplant Analysis

### 5.1. Background

Liver transplant surgery is the only life-saving treatment for patients in certain medical conditions. It is a challenging surgical procedure, and significant medical resource, experience and dedication are needed. During the surgery, the clamping of major vessels and the subsequent vascular anastomosis bring huge impacts on the recipient’s circulation system [39]. As a better understanding of the cardiovascular dynamics during the procedure may help optimize the intraoperative management, commercial monitoring instruments based on real-time arterial blood pressure (ABP) waveform analysis has been introduced. However, they been questioned subsequently for its performance in liver transplant surgery [5, 46, 42]. Thus, obtaining useful information from the ABP waveform in liver transplant is still a challenging problem.

Traditionally, various extracted features, either landmark measurements in the time domain [33] or quantities in the frequency domain [49], serve as the input for the subsequent pulse waveform analysis. These designated features are supposed to reflect underlying physiological information, or those parameters driving the network interaction. However, it is reasonable to suspect that information hidden in the finer scale might be ignored via the above approach, and hence finer structure of network dynamics is overlooked, particularly when the physiology is disturbed. It is thus reasonable to argue that taking the whole waveform into account might provide more complimentary information compared with those traditional parameters. On the other hand, due to the short-term memory nature of human brain, it is challenging to visualize and directly utilize the dynamics encoded in the ABP waveform on the large scale. Motivated by handling the above challenges, including finding finer information on the short scale, and exploring the dynamics on the large scale, in our previous research, we reported a solution under the manifold learning framework, and showed that the DM can extract rich information directly from the *raw cardiovascular waveform* [48]. The novelty in [48] capturing subtle morphological changes that might be overlooked by the designed features. However, due to the computational barrier intrinsic to the DM, the approach is limited to relatively small dataset.

In this study, we hypothesize that with the help of Roseland, the manifold learning approach shown in [48] can be applied to study the ABP waveform during the liver transplant procedure, and provide hemodynamic information on both the small and large scales. Note that the Roseland is critical in this analysis since the whole period of the surgery can yield more than 10^5^ continuous pulses as data points in high dimensional space for a pulse-to-pulse waveform analysis.

### 5.2. Material

The data was collected from an observational study per institutional ethic regulation. We collected physiological signals via the data collection software, S5 collect (GE Healthcare, Chicago, Illinois, United States) from the standard patient monitor instrument, GE CARESCAPE^TM^B850 (GE Healthcare, Chicago, Illinois, United States). The recorded ABP signal was uniformly sampled at 300 Hz in the instrument and resampled at 500 Hz via the cubic spline interpolation for off-line processing. The signal is of 78,350s long spanning the whole surgical procedure and contains 120,725 pulses.

### 5.3. Data analysis

Denote the ABP waveform as *x*^*A*^ ∈ ℝ^*N*^. We used the maximum of the first derivation during the ascent of each ABP pulse waveform as a fiducial point. A legitimate ABP pulse is determined by a two-pass algorithm using the following measurements automatically: the peak maximum, the trough minimum, the minimum of difference between the maximum and minimum within the pulse, the pulse width, and the duration to the previous pulse. The thresholds for those measurements are automatically adjusted by a feedback mechanism. Suppose there are *L* legitimate cycles in *x*^*A*^. Denote the *i*-th fiducial point as *n*_*i*_. Break *x*^*A*^ into *L* − 1 segments so that the *i*-th segment is the *i*-th ABP pulse containing one waveform cycle. Denote the *i*-th segment as 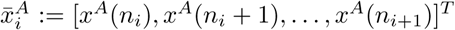. Since the duration of each pulse is not constant, we truncated them into an uniform size according to their minimal length *q* = min *n*_*i*+1_ *n*_*i*_ +1 ℕ, and get 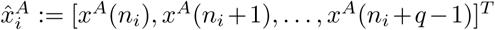. Next, normalize 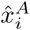 by removing the mean and setting the variance to 1 to separate the blood pressure information from the normalized ABP pulse, and denote the normalized ABP pulse as 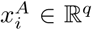. Derived from the ABP signal, we get the data set 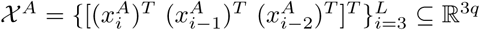. We assume that 𝒳 ^*A*^ can be well approximated by a low dimensional manifold, referred to as the *wave-shape manifold* [28]. To apply the Roseland, the landmark set 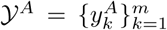, where *m* = ⌊*n/*600⌋, was chosen from 𝒳 ^*A*^ via setting 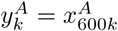.

### 5.4. Results

The dataset consists of *L* = 120, 725 legitimate cycles of length *q* = 334, and hence 120, 723 data points in 𝒳^*A*^. Thus, *m* = 201 and note that 201 = 120, 723^0.453^. The total computation time of the Roseland algorithm is less than 40s on an ordinary personal computer (CPU: Intel Core i5-7500, operation system: Microsoft Windows 10 64-bit home edition, programming platform and language: Microsoft Visual Studio Community version 2019, .NET framework 4.8, and C#, LAPACK software library: Intel Math Kernel Library 2020 Initial Release), while the estimated computation time using traditional DM algorithm based on eigendecomposition would be more than one day.

The embedding result is shown in Figure 6. The successive pulses evolve with time and constitute a trajectory on the manifold presented as a 3D embedding (Fig.6, panel A). The trajectory visits different locations during different steps of the liver transplant procedure. Moreover, there is a “clustering” effect in the embedding, which is enhanced by the imposed color that encodes the temporal information. We can thus visualize the relationship among different hemodynamic status during different surgery steps. This relationship provides physiological dynamics on the large scale. We emphasize that while we can easily read the waveform, but it is not easy to perceive the dynamics and organize them with only human eyes and brain (Fig.6, panel B and C), particularly when the signal is long.

**Figure 6.**
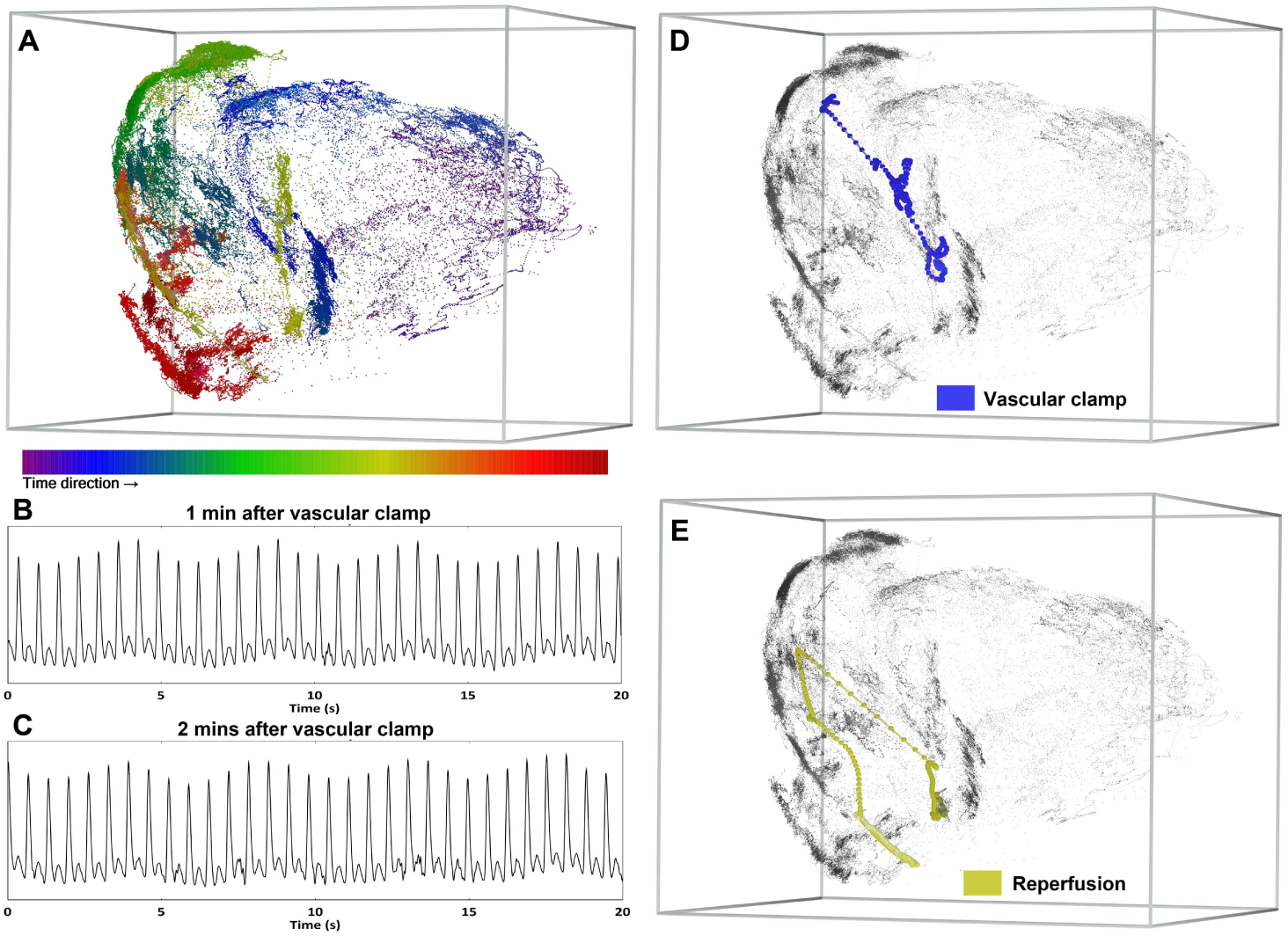
The 3D embedding (panel A) of pulse-to-pulse pressure waveforms collected from a 78,350s ABP signal (120,725 pulses) during the liver transplantation procedure. The embedded pulses are labeled by colors encoding the time. The color helps visualize the ever-changing trajectory formed from successive pulse waveforms. The embedding is clustered and different clusters are related to different stages. During transition phases of inferior vena cava cross clamping (panel B, C), the ABP tracings provide little clues with respect to the subtle waveform information and its longterm evolving, while the Roseland algorithm reveals the fast paced movements (panel D). To signify the physiological dynamics associated with the vascular clamp (panel D) and reperfusion (panel E) events, pulses in transition phases are labeled with colored linked dots while the rest pulses are not colored. In panels A, D, and E, the grids are drawn to enhance the 3D visualization, and an online supplementary video is provided for more details.

We further quantify the trajectory in different surgical phases as well as the phase transition periods in which the trajectory moves in fast pace. We consider the following different hemodynamic phases during the liver transplant procedure, particularly those that phase transition happens with violent physiological changes take place—First, the occlusion of blood inflow to the “old” liver (to be replaced), performed by the cross-clamp of the inferior vena cava, the largest vein of the human body; second, the start of the circulatory connection from the graft (the new liver organ) to the circulation system as the blood flow starts in the portal vein; third, the start of the connection between hepatic artery and the graft. All these transitions drastically affect the cardiovascular system via the changes of fluid volume and electrolyte composition.

For each hemodynamic phase, we embed the ABP waveforms by the Roseland embedding into 10-dim Euclidean space; that is, *q*′ = 10, and evaluate the geometric center of all beats during that phase. Then, we define the distance between two hemodynamic phases groups by measuring the RDD between their geometric centers. The quantitative measurement is expressed as mean and 95 % confidence interval after bootstrap resampling without replacement in 100,000 samples. In light of the Newtonian mechanics, we consider two quantities. The *two-point velocity* measures the dynamics on the large scale. The two-point velocity is defined as the ratio of the RDD between two points and the time difference between the two points. We also consider the *trajectory speed*, which measures the dynamics on the small scale. The trajectory speed is defined as the ratio of the path length of the trajectory and the time different between the beginning and ending of the trajectory. The quantitative result (Table 1) shows that during the phase transition, the trajectory moves faster macroscopically (on the large scale) when quantified by the two-point velocity, particularly when compared with that within each surgical phase. However, the trajectory speeds, which represents hemodynamics on the microscopic scale (small scale), are similar during the phase transition and within each surgical phase. The numeric results are consistent with visualization from the 3D embedding (Figure 6 and 7).

**Table 1.**
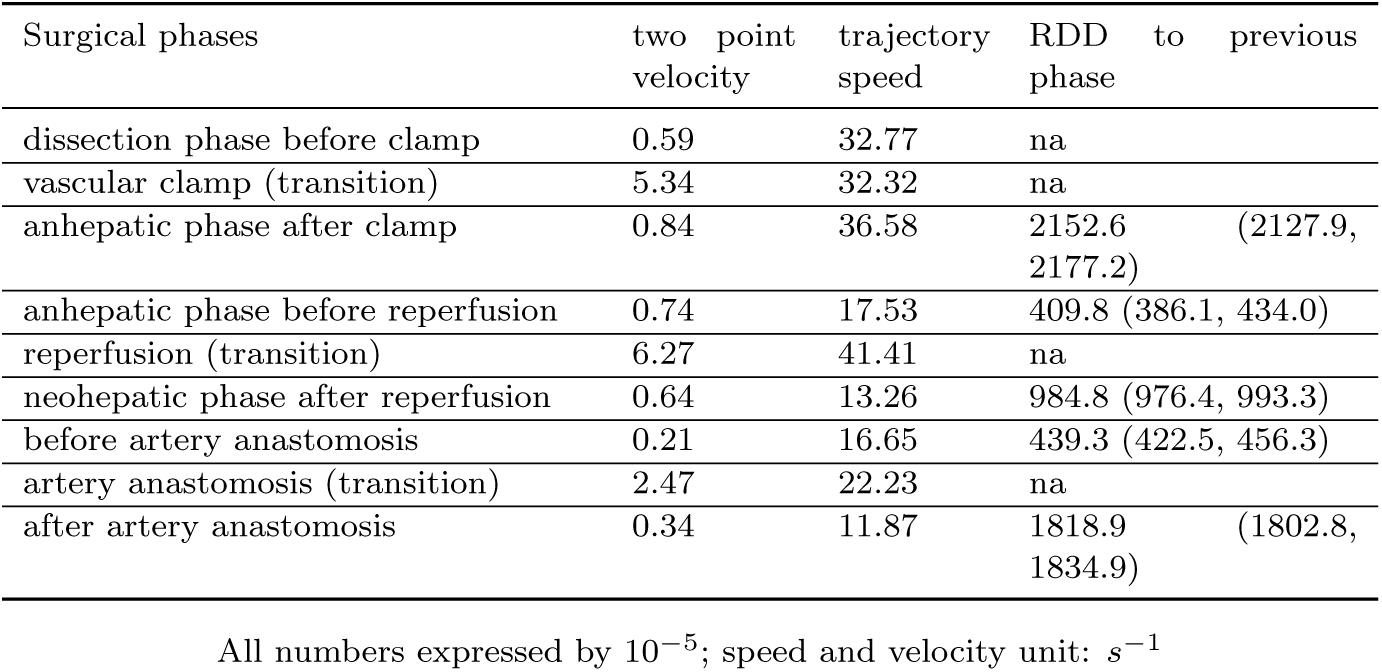
Quantitative results of phases and phase transitions from liver transplantation data

| Surgical phases | two point<br>velocity | trajectory<br>speed | RDD to<br>previous<br>phase |
| --- | --- | --- | --- |
| dissection phase before clamp | 0.59 | 32.77 | na |
| vascular clamp (transition) | 5.34 | 32.32 | na |
| anhepatic phase after clamp | 0.84 | 36.58 | 2152.6 (2127.9,<br>2177.2) |
| anhepatic phase before reperfusion | 0.74 | 17.53 | 409.8 (386.1, 434.0) |
| reperfusion (transition) | 6.27 | 41.41 | na |
| neohepatic phase after reperfusion | 0.64 | 13.26 | 984.8 (976.4, 993.3) |
| before artery anastomosis | 0.21 | 16.65 | 439.3 (422.5, 456.3) |
| artery anastomosis (transition) | 2.47 | 22.23 | na |
| after artery anastomosis | 0.34 | 11.87 | 1818.9 (1802.8,<br>1834.9) |

All numbers expressed by $10^{-5}$ ; speed and velocity unit: $s^{-1}$

**Figure 7.**
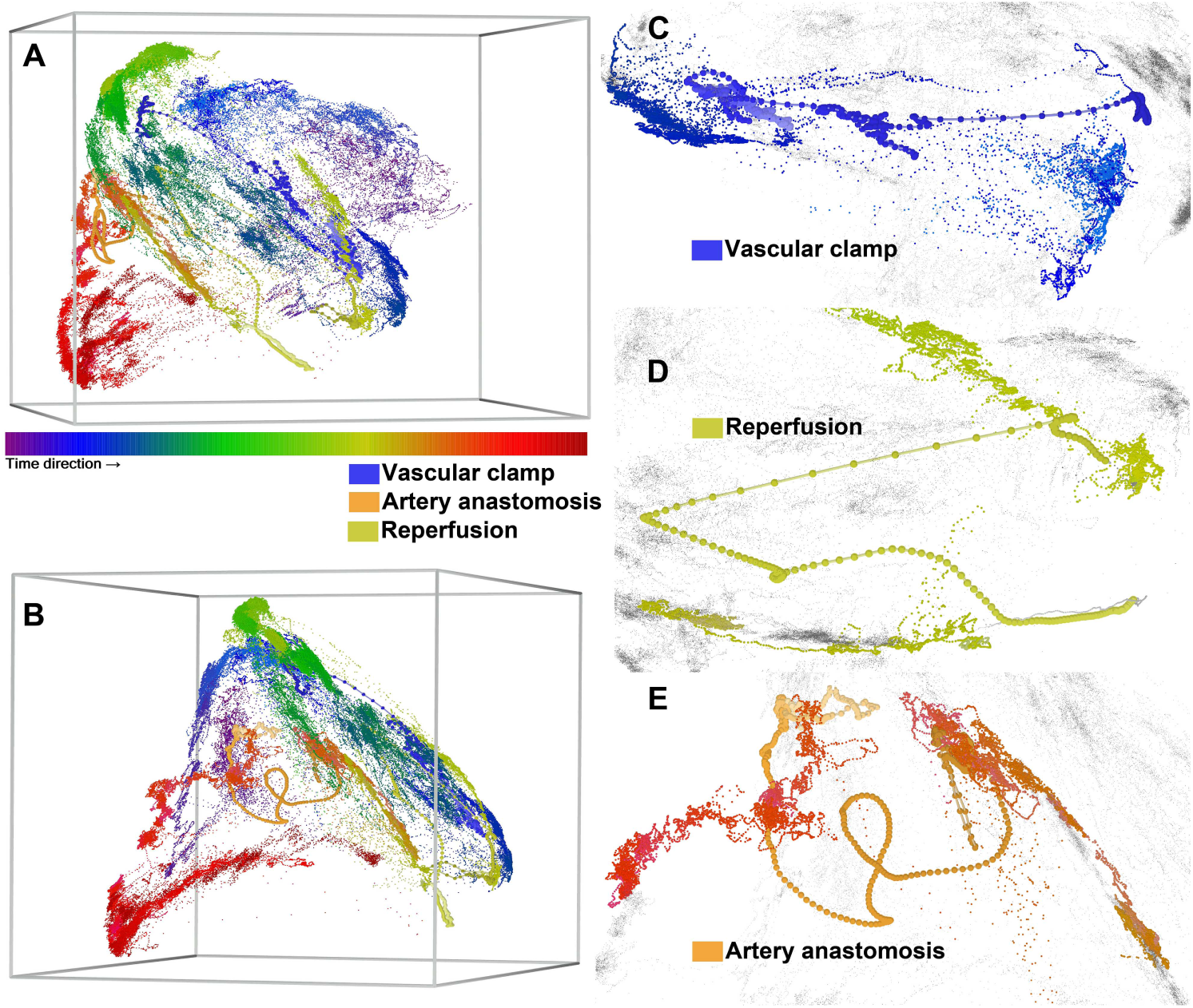
The 3D embedding highlights the locations of transition phases (in linked dot) in relation to the whole period of ABP waveforms (smaller dot) in a liver transplant surgery. The color labels the time sequence. Panel B is the horizontally rotated Panel A by 60 degrees for a better visualization of the artery anastomosis transition. These views show geographic relationship between surgical stages and transition phases. Zoom-in views of the 3D embedding of pulse-to-pulse ABP waveform include transition phases of major vascular cross-clamping (panel A), new liver graft reperfusion (panel B), and hepatic artery anastomosis (panel C), which shows fast paced movement in transition phases (enlarged colored dots with line-link). On the other hand, in the statuses immediate before and after transition (colored dots without line-link), we see less movement and the embeddings are clustered. The rest pulses (uncolored small dots) appears in the background.

The liver-transplant example shows the benefit and potential of the Roseland algorithm. As more (longer) data leads to a richer knowledge base, we now have an unprecedented signal processing tool for a long-period signal with complex underlying physiology. In the liver transplant example, the 3D embedding reflects the complex relationship among different surgical phases without *ad hoc* pulse waveform knowledge. This suggests the practicability of handling data governed by complex physiological mechanism. As there is room to be improved in monitoring the hemodynamic status in liver transplant surgery [5, 46, 42], we expect that the proposed waveform analysis would lead to more insights into the hemodynamic status of liver transplant surgery to improve the patient’s outcome. We will report research in this direction in our future work. Certainly, the source of knowledge base is not limited to the ABP waveform. Different physiological waveforms can be considered to further enrich the knowledge base. How to simultaneously utilize multimodal physiological waveforms, particularly when the recording is long, is a relatively white area, and we expect that the proposed waveform analysis would form a base toward this goal. We also expect that the similar principal could be applied to study other medical datasets for different medical problems, for example, the long-term outcome of the patient underwent organ transplantation with respect to the immune function, or the genetic predisposition and environment factors with respect to the cancer occurrence.

## 6. Discussion

### 6.1. More related work – scalability and robustness

The recent paper [10] contains a comprehensive review of numerical acceleration techniques for nonlinear dimension reduction, and we refer readers with interest to that work. To handle scalability, an intuitive approach is accelerating the kNN search step. See [10] for a summary and a recently proposed randomized kNN approach [29]. However, it is well known that the kNN scheme is not robust when the dataset is noisy when the neighboring information is not provided. Specifically, it is challenging to estimate pairwise distance robustly, unless we have extra structure to design a robust metric, for example, in the image analysis [8]. If the tangent plane is known, it can help us determine neighbors [47]; however, when the dataset is noisy, the local principle component analysis approach to estimate the tangent space is biased [21]. In short, the kNN is only useful when we have an accurate information about the neighbors.

Another natural approach to handle scalability is accelerating the eigen-decomposition step. For example, we can approximate the kernel decomposition by classical iteration-based algorithms [18]. We can also evaluate the matrix decomposition by designing a randomized algorithm [37].

For the robustness issue, one naive idea is “denoising” the dataset before applying any algorithm. However, it is in general an independent challenging problem. Under the manifold setup, researchers have proposed several algorithms to denoise the dataset. For example, the “reverse diffusion” scheme [20] and the manifold fitting scheme [13]. We mention that the algorithm might not be scalable, but not too much is known at this moment. Another approach is modifying the random walk scheme to a non-lazy random walk via diffusion to obtain a self-consistency Markov chain. But it is under the assumption that the edge information is known [44], which is not possible in many applications. To our knowledge, the general theory for the robustness of kernel methods was first studied in [11], and the analysis was extended to the large noise setup [12]. The authors proved that the spectral embedding methods can be efficiently stabilized by forcing the random walk to be a non-lazy one on the complete graph. Unfortunately, while it could help stabilize the noise impact, the algorithm is not scalable.

### 6.2. Relationship with the alternating diffusion algorithms

Note that the Roseland is related to the recently developed alternating DM (ADM) algorithm [24]. The ADM is developed to deal with the sensor fusion problem, when we have multiple data-sets simultaneously acquired by multimodal sensors. In short, suppose we have 2 aligned data sets 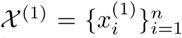 and 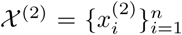, then we build two transition matrices *K*^(*j*)^ = *D*^−1^*W* from 𝒳^(*j*)^, where *j* = 1, 2. Next the alternating diffusion operator is defined as *K*^(1)^*K*^(2)^. It is easy to check that *K*^(1)^*K*^(2)^ is also row stochastic, so it can be considered as a transition probability matrix of a new Markov chain that alternates between the two data sets. The idea of the Roseland is closely related to the ADM, in the sense that we can consider the landmark set as a new data set on its own right. So, the idea of measuring similarity between two data points via the landmark set can be understood as diffusing between the original data set and the landmark set from the ADM point of view.

### 6.3. Application of the Roseland

The idea of landmark set have several applications. Here we mention two of them. The VDM [43] is a generalization of DM that aims to encode the group structure when comparing objects. The VDM suffers from the expensive computational cost more than the DM, since the group structure is usually represented as a matrix, which inflates the matrix size. Specifically, if the group structure is represented as a *q* × *q* matrix and we have *n* objects to compare, then we need to eigendecompose a *nq* × *nq* kernel matrix in the VDM. We expect the landmark idea can be generalized to accelerate the VDM. We will explore this possibility in our future work.

Spectral clustering methods are known to perform well when the classical clustering methods such as *k*-means and linkage fail [1]. It is well known that the more clusters we need to determine, the more eigenvectors we need [1, 25]. As is shown in the numerical section, the Roseland has the ability to recover more and better eigenvectors, at least compared with the Nystöm extension. This shows the potential of applying the Roseland for the multiway spectral clustering purpose.

## Acknowledgement

The authors acknowledge Dr. Shen-Chih Wang for the fruitful discussion and suggestion. The work of Yu-Ting Lin was supported by the National Science and Technology Development Fund (MOST 108-2115-M-075-001) of Ministry of Science and Technology, Taipei, Taiwan.

